# Rehabilitation drives functional reorganization of intact corticospinal-supraspinal projections following partial spinal cord injury

**DOI:** 10.64898/2025.12.12.694014

**Authors:** James Bonanno, Sheel Trivedi, Ciara F. O’Brien, Sharna Saha, William B.J. Cafferty

**Author notes:** Corresponding Author: William B.J. Cafferty, Department of Neurology, Yale University School, New Haven, Connecticut 06511. **Present Address (if applicable)**: Cold Spring Harbor Laboratory School of Biological Sciences, 1 Bungtown Road, Cold Spring Harbor, New York 11724 (Ciara F. O’Brien).

## Abstract

Spinal cord injury (SCI) disrupts corticospinal tract (CST) connectivity and impairs skilled voluntary movement. However, most human SCIs are anatomically incomplete, allowing spared CST pathways to engage in rehabilitation-mediated plasticity to promote functional recovery. How voluntary rehabilitation engages and reorganizes the supraspinal targets of the intact CST remains incompletely understood. Here, we combined unilateral pyramidotomy (uPyX) in male and female mice with continuous voluntary complex-wheel running to test whether fine motor-dependent rehabilitation drives supraspinal CST plasticity. uPyX mice rapidly resumed wheel running after a transient deficit. In contrast to lesion-only controls, rehabilitation significantly improved skilled forelimb performance on the horizontal ladder rung task. Immunohistochemical c-Fos labeling confirmed that complex-wheel running robustly activated the intact forelimb CST in motor cortex. Whole-brain CST projection mapping using intersectional viral vector tracing revealed targeted supraspinal reorganization localized to medullary motor nuclei. Three nuclei – the lateral paragigantocellular reticular nucleus (LPGi), gigantocellular reticular nucleus, alpha part (GiA), and ventral medullary reticular nucleus (MdV) – exhibited significant lesion– and/or rehabilitation-induced increases in CST innervation. Rehabilitation-driven CST sprouting correlated with regional c-Fos activation, indicating activity-dependent remodeling. Notably, CST projection density in the MdV, critical for skilled forelimb control, correlated with functional recovery. These findings identify a set of spinally-projecting medullary nuclei as key sites of rehabilitation-induced CST plasticity and highlight the MdV as a potential mediator of restored motor function. This work defines how voluntary rehabilitation reorganizes spared corticospinal pathways and provides targets for optimizing activity-based interventions after SCI.

**Significance Statement:** Effective rehabilitation after spinal cord injury (SCI) must harness the plasticity of spared motor pathways, yet the supraspinal circuits that support rehabilitation-mediated recovery remain unknown. Using a model that preserves voluntary motor engagement, we show that continuous fine motor-dependent rehabilitation activates intact corticospinal neurons and drives highly specific remodeling of their supraspinal terminals. Rehabilitation selectively strengthens CST inputs to motor regions of the medulla, particularly the ventral medullary reticular nucleus (MdV), and CST plasticity within this region predicts enhanced behavioral recovery. These findings highlight the MdV as a central locus by which rehabilitation re-establishes descending control of the impaired limb, providing mechanistic insight to guide targeted, circuit-based rehabilitation therapies for incomplete SCI.

## Introduction

Spinal cord injury (SCI) causes devastating and often permanent loss of voluntary and fine motor control (Mensah et al., 2025). However, most human SCIs are anatomically incomplete, leaving spared motor pathways that can support a limited degree of spontaneous functional recovery (Raineteau and Schwab, 2001; Burns et al., 2012). Currently, intensive rehabilitation is the only established means of enhancing functional recovery in both humans (Lee and Jeoung, 2023) and animal models (Battistuzzo et al., 2012), and emerging evidence indicates that combining it with neuromodulation may further engage spared circuits and strengthen activity-dependent plasticity (Asboth et al., 2018; Wagner et al., 2018). Because adult nervous system axons are unable to regenerate (Liu et al., 2006), rehabilitation is thought to promote functional recovery through neuroplasticity of spared circuits in the brain and spinal cord (Loy and Bareyre, 2019), but the exact anatomical substrates that enable rehabilitation-induced recovery are still poorly understood.

One key target for understanding rehabilitation-induced functional recovery is the corticospinal tract (CST). Located across layer 5 of the sensorimotor cortex, the CST is the primary descending motor pathway that drives voluntary, fine-motor control in mammals (Iwaniuk and Whishaw, 2000; Lemon and Griffiths, 2005). Evidence strongly demonstrates that the intact motor cortex gains new functional control over affected limbs in animal models of partial SCI (Ghosh et al., 2009; Carmel et al., 2014) and stroke (Lindau et al., 2014). Additionally, in animal models of partial SCI (Weidner et al., 2001; Oudega and Perez, 2012), unilateral stroke (LaPash Daniels et al., 2009), and TBI (Dancause et al., 2005), the CST exhibits a capacity for sprouting in the spinal cord and supraspinal motor regions. This sprouting can be enhanced by acute electrical stimulation of the motor cortex (Brus-Ramer et al., 2007), genetic manipulations (Cafferty and Strittmatter, 2006; Lindau et al., 2014), and motor rehabilitation (Starkey et al., 2011; Liu et al., 2016). Moreover, supraspinal motor cortex activation when coordinated with spinal activation leads to enhanced motor outcomes after injury (Van Steenbergen et al., 2022). Together, these studies suggest that activity-dependent plasticity of the CST, driven by rehabilitation, can be leveraged to improve functional recovery from SCI.

Recently, data from our laboratory (Golan et al., 2023) and others (Nelson et al., 2021; Carmona et al., 2024) have highlighted and characterized the breadth of CST-supraspinal projections. Specifically, we have identified that the striatum, thalamus, midbrain, pons, and medulla receive input from the CST. While the ventral medulla has previously been identified as a region that undergoes rehabilitation-induced plasticity from the CST (Asboth et al., 2018), a brain-wide investigation of how intact CST-supraspinal terminals reorganize after injury and rehabilitation has never been performed.

Here, we used a unilateral pyramidotomy (uPyX) model combined with continuous voluntary fine motor rehabilitation, whole-brain projection and c-Fos mapping to test whether rehabilitation drives supraspinal remodeling of the intact CST. We found that rehabilitation enhances skilled motor recovery and increases lesion-induced sprouting in motor regions of the medulla. Crucially, we found that intact CST projections in spinally-projecting regions of the motor medulla reorganize in rehabilitated animals in an activity-dependent manner. Finally, we identified three nuclei of the motor medulla that undergo rehabilitation-induced sprouting and found that sprouting in the ventral part of the medullary reticular formation – a key region for fine forelimb function (Esposito et al., 2014) – correlated with improved functional recovery. These insights into the circuit mechanisms by which voluntary rehabilitation reorganizes spared corticospinal pathways lay the groundwork for developing targeted interventions that amplify rehabilitation-driven plasticity and improve motor outcomes following central nervous system injury.

## Materials and Methods

### Mice

C57BL/6 mice (The Jackson Laboratory, Charles River Laboratory) were used for all behavioral and anatomical experiments. The mice were maintained in a temperature and humidity-controlled room on a 12 h light/dark cycle with lights on from 7:00 A.M. to 7:00 P.M with *ad-libitum* access to food and water. For experiments, mice were randomly assigned to groups.

### Surgery

All procedures and postoperative care were performed in accordance with the guidelines of the Institutional Animal Use and Care Committee at Yale University.

*Unilateral Pyramidotomy (uPyX):* Mice received a left side unilateral pyramidotomy in the medulla rostral to the pyramidal decussation as previously described (Cafferty & Strittmatter, 2006). Briefly, mice were anesthetized with ketamine (100 mg/kg; Covetrus) and xylazine (15 mg/kg; Covetrus) and placed in a supine position, an incision was made to the left of the trachea, and blunt dissection exposed the occipital bone at the base of the skull. Depth of anesthesia was continuously monitored via reflexes and depth of breath. The occipital bone was removed on the left side of the basilar artery with blunt Dumont #2 forceps to expose the medullary pyramids. The dura mater was pierced with a 30-gauge needle and resected. The left pyramid was transected unilaterally with fine Dumont #5 forceps to a depth of 0.25 mm or exposed for sham lesion. No internal sutures were made, and skin was closed with monofilament suture. All animals received pre-surgical analgesia (Buprenorphine extended release 3.25 mg/kg subcutaneously), and post-surgical antibiotics (Ampicillin, 100 mg/kg subcutaneously) for 2 days post lesion. Histological verification of lesion completeness was performed after the completion of behavioral experiments using PKC gamma staining of cervical spinal sections (Cell Signaling Technology, #59090).

*Intersectional CSN Labeling:* Intersectional viral tracing was completed at day 28 post lesion to ensure no tracing dependent functional deficits contributed to behavioral assessments. To label forelimb CSNs, adult mice were anesthetized with ketamine (100 mg/kg; Covetrus) and xylazine (15 mg/kg; Covetrus) and placed in a stereotaxic frame (Stoelting). First, a small incision was made into the skin and retracted using hemostats. Small craniotomies were then made over the centers of mouse rostral forelimb area (RFA), coordinates +2.0-2.5 mm AP and +0.5-1.5 mm ML from bregma, and caudal forelimb area (CFA), coordinates –0.5-1.0 mm AP and +1.0-1.5 mm ML from bregma. The tip of a pulled glass capillary tube attached to a 5 µl Hamilton syringe was loaded into a Micro4 infusion device (World Precision Instruments) for a total of two injections of pAAV1-CAG-FLEX-eGFP (Addgene, #59331) into RFA and three injections of pAAV1-FLEX-tdTomato (Addgene, #28306) into CFA at a depth of 500 µm. Each injection site received a total of 100 nL of virus at a rate of 40 nL/min. After each injection, the capillary tube was kept in place for 30 sec before being pulled up slowly. After all injections were completed, the skin was sutured with 4.0 Vicryl (McKesson) and mice prepared for spinal injections.

During the same procedure, an incision was made over the cervical enlargement to reveal C5-C8 vertebrae by first dissecting overlying muscle. Then, a unilateral left side laminectomy was performed to expose the underlying spinal cord. The same injection apparatus was used for spinal injections. For each spinal level in the C5-C8 range, the capillary tube was slowly inserted stereotaxically to a depth of 500 µm into the spinal cord and ∼600 µm lateral from the midline. 30 seconds after the introduction of the capillary tube, 100 nL of pENN-rAAV-.hSyn-Cre-WPRE-hGH (Addgene, catalog #105553) was infused into the spinal cord over 2 min. The tip was left *in situ* for an additional 30 sec before removal. After all injections, the skin was sutured with 4.0 Vicryl, and mice were monitored until they awoke from anesthesia. All animals received pre-surgical analgesia (Buprenorphine extended release 3.25 mg/kg subcutaneously), and post-surgical antibiotics (Ampicillin, 100 mg/kg subcutaneously) for 2 days post lesion.

*Voluntary Complex Wheel Running Rehabilitation:* Voluntary rehabilitation was administered to mice (2–3 months old, *n* = 9 males, 13 females) using our previously described REVS hardware and software system (Bonanno et al., 2025). Briefly, mice randomly assigned to rehabilitation groups were given access to running wheels for five days prior to lesion for habituation. This included two days of regular wheels, followed by three days of complex wheels. Regular wheels had 1 cm rung separations and complex wheels had a mix of 1-3 cm rung separations. Mice had access to running wheels 24 hours a day, except for the 48 hours immediately after the lesion/sham surgery. Mice (n=1, uPyX+Rehab) that failed to engage in wheel running were removed from the study and excluded from analyses. Wheels were cleaned with water and ethanol once a week after ladder rung testing.

*Complex Ladder Rung:* Mice (2–3 months old, *n* = 27 males, 18 females) were trained for forepaw and hind paw dexterity assessment using an irregular horizontal ladder. Mice were handled for 3 days prior to any behavioral testing and were allowed to habituate to the ladder apparatus with regularly spaced rungs for 5 minutes. After habituation, mice were trained to cross the regular ladder to a safe box with bedding. A bright light was placed at the start of the ladder to encourage crossing to the safe box. Experimental handling was carried out by an experimenter blinded to lesion status. A GoPro Hero8, set up 7.5 inches from the apparatus, was used to capture video recordings of performance on the irregular ladder rung (Metz and Whishaw, 2009) at baseline and 2-, 7-, 14-, 21-, and 28-days post lesion. Rungs were spaced 1, 2, or 3 cm apart in a randomized pattern which was changed between testing sessions to prevent learning. In an individual behavior session, mice crossed the irregular ladder three times, resulting in a total of 1 meter of distance traveled.

To analyze paw placement as an injury metric, first DeepLabCut (Mathis et al., 2018) was used to track performance of the forepaws and hind paws for each mouse. A network was trained, using 300 labeled frames selected from a random mix of sham and lesioned animals across all days. The network was refined until predictions were less than 5 pixels off compared to human-labeled frames. A total of 166 videos were analyzed using the network. Videos were manually preprocessed for DeepLabCut by cutting out frames without the mouse. Output CSV files were processed and analyzed using a custom-written peak detection algorithm in R to detect missed steps using paw y-position. A cut-off was set to count any placement of the paw below the center of each rung as a missed step. Total number of missed steps calculated for each limb. Injured forelimb reach errors, or overreaches, were quantified by setting a threshold two standard deviations above the mean step height. Time to cross was calculated by multiplying total frames by the frame rate of the camera.

To create paw trajectory plots, individual reach and footfall values were isolated using custom-written analysis code described above. We plotted individual successful reaches and footfall observations for the baseline, 2 dpi, and 28 dpi conditions. To visualize a mean or “typical” reach and footfall, we applied locally weighted polynomial smoothing (LOESS), a nonparametric regression method implemented in the ggplot2 library in R. Shaded ribbons indicate 99% confidence intervals around the LOESS-estimated mean trajectory.

*Immunohistochemistry:* Ten days after intersectional AAV injections, mice were euthanized with isoflurane and transcardially perfused with 0.9% NaCl with 10 units/ml heparin (Covetrus, 1000 unit/ml) followed by 4% paraformaldehyde (PFA) in PBS. Brains and spinal cords were dissected and post-fixed in 4% PFA overnight at 4°C. The next day, 75 µm thick sections of brain and 35 µm of cervical spinal cord were cut using a vibratome (Leica Microsystems, VT1000S), and sections prepared for free-floating immunohistochemistry.

*Corticospinal*-*supraspinal terminal detection and c-Fos activation*: Every other section throughout the brain was stained using free floating IHC. Sections were washed in 0.2% PBS-T 3x and blocked in 10% Normal Donkey (NDS) or Normal Goat Serum (NGS) for 1 hour. Primary antibodies to enhance RFP and GFP signal, Rabbit anti-RFP (Abcam; ab13970, RRID:AB_300798) and Chicken anti-GFP were used (Abcam; ab167453, RRID:AB_2571870) in addition to Guinea Pig anti-c-Fos (Synaptic Systems, 226 308, RRID:AB_2905595) were diluted 1:1,000 in PBS-T in 10% serum overnight at RT. The following day, sections were washed 3x in PBS-T, and incubated in secondary antibodies (Alexa Fluor 546: A10040, A32732; Alexa Fluor 488: A78948, A32931; Alexa Fluor 647: A21450, ThermoFisher) for 2 hrs. Next, sections were washed 3x in PBS-T. The final wash contained a 1:10,000 dilution of 1 mg/ml DAPI stock solution (ThermoFisher, 62448). Sections were then mounted onto Superfrost Plus slides and treated with TrueBlack (Biotium, 23007) for 30 secs diluted 20x in 70% EtOH to reduce autofluorescence and background staining. Slides were then washed 3x in PBS and cover slipped with Fluoro-Gel (Fisher Scientific, 17985-11). Whole-section imaging at 4x was performed using an Olympus VS200 Slide Scanner.

*PKC-γ detection for lesion completeness*: Histological verification of lesion completeness was performed after the completion of behavioral experiments using PKC-γ staining of cervical spinal sections. Cervical spinal sections were washed in 0.2% PBS-T 3x and blocked in 10% Normal Donkey (NDS) for 1 hour. Sections were then incubated with Rabbit anti-PKC-γ (Cell Signaling Technology, #59090) primary antibody overnight at RT. The following day, sections were washed 3x in PBS-T and incubated in Donkey anti-rabbit Alexa Fluor 647 (A31573, Fisher Scientific) secondary antibodies for 2 hours, then washed 3x in PBS-T and mounted with Fluoro-Gel (Fisher Scientific, 17985-11). Mice were excluded (n=4 uPyX+Rehab, 4 uPyX-Rehab) from the study if lesioned side ventral dorsal column staining was assessed to be 5% or more intact.

*BrainJ Projection Density Analysis:* BrainJ software was used to label CSN soma and axonal projections throughout each brain, as previously described (Botta et al., 2020). Briefly, sections were manually ordered and registered to the Unified Kim Mouse Brain Atlas (Chon et al., 2019) template using Elastix software, version 5.0.1. Next, ilastik, version 1.3.3 was used to identify brainwide soma and projections in the ∼75 mounted sections using pixel-based machine learning. Ilastik training was performed by two experimenters blinded to groups. BrainJ output absolute density measurements for 1429 subregions of the brain, as defined by the Kim Atlas. Projection density for each animal was normalized to the total projection density measured in the corticospinal fiber tracts. For higher level projection analysis, we assigned subregions to one of fifteen functional regions: isocortex, striatum, thalamus (sensory-motor), thalamus (other), midbrain (motor), midbrain (sensory), midbrain (other), pons (motor), pons (sensory), pons (other), medulla (motor), medulla (sensory), medulla (other), and cerebellar nuclei.

### Experimental Design and Statistical Analysis

*Behavioral experiments*: There were four groups including both sexes: Sham without rehabilitation (Sham-Rehab, n=8), Sham with rehabilitation (Sham+Rehab, n=7), uPyX without rehabilitation (uPyX-Rehab, n=11), and uPyX with rehabilitation (uPyX+Rehab, n=11). Rehabilitation metrics were collected from rehabilitation groups only using REVS, and the metrics analyzed included total distance, total hours run, and overall speed. Cumulative distance run was separately calculated by summing the total distance per night for each animal. Each rehabilitation metric was evaluated using a mixed-model ANOVA with a between-subjects variable of lesion status and a within-subjects variable of day. Post-hoc t-test comparison of baseline and 2 days post-injury (dpi) specifically were performed for each metric to assess acute deficits.

For the complex ladder rung task, total injured forelimb steps, injured forelimb footfalls, intact forelimb footfalls, injured forelimb reach errors, injured hindlimb footfalls, and intact hindlimb footfalls were measured to assess fine motor control, and total time to cross was measured as a general metric of impairment. Complex ladder rung metric was assessed using mixed-model ANOVAs with between-subjects variables of lesion status and treatment (rehabilitation) and a within-subjects variable of day. Separate ANOVAs were run for the baseline to 2 dpi time period and the 7 dpi and 28 dpi time periods to isolate the effects of lesion and rehabilitation. When significant main effects were detected, post hoc t-tests were performed to identify specific differences between groups on specific days. Post hoc testing was corrected for multiple comparisons using Bonferroni corrections. A principal component analysis (PCA) was performed on injured forelimb steps, footfalls, reach errors, intact footfalls, injured hindlimb footfalls, intact hindlimb footfalls, and time to cross across –2, 2, and 28.

*Colocalization analyses*: CSNs and c-Fos positive cells were labeled using ilastik. A list of detected CSNs and c-Fos positive cells, and all channel intensity values for each cell, was output for each brain. The threshold for a CSN to be classified as “positive” was the minimum intensity value of c-Fos detected cells for a given brain. For analyses, CSNs were automatically assigned to M2 and M1 of motor cortex based on the Kim Atlas, and measurements from RFA– and CFA-traced CSNs were included. CSNs assigned to layers 1 and 6 were removed from analyses. Linear mixed-effect models were performed with colocalization proportion as the dependent variable, lesion status and treatment as fixed effects, and subject ID as a random intercept. A subset (n = 12) of mice in the rehabilitation and complex ladder rung experiments received c-Fos staining. For this experiment Sham+Rehab and uPyX+Rehab mice were confirmed to have run within the previous hour using REVS.

*Projection Density Analyses*: The dependent variable was projection density in a given region, as defined by total signal occupied divided by total region area. Projection density measurements were normalized for each animal (n = 9 control, 5 uPyX-Rehab, 5 uPyX+Rehab) based on the total projection density measured in corticospinal fiber tracts, as measured by BrainJ. Mixed-effect linear models were used for both the higher level (i.e., medulla-motor) and granular (i.e., MdV) projection density analyses. The Allen Mouse Brain CCF does not include a paramedian reticular nucleus; therefore, PMn and MdV measurements were combined and designated as MdV to facilitate comparison with Allen-based anatomical designations. Atlas Mice with failed tracing and incomplete hindbrain collection were excluded from analyses. Lesion status and treatment (rehabilitation) were used as fixed effects in the model, with animal ID included as a random effect. For sub-region analysis, we restricted our analyses to spinally-projecting subregions, as identified by Wang et al., 2022. Post-hoc t-tests were only performed on subregions identified as having significant interactions between region and group. P-values were corrected for multiple comparisons using the Tukey method for adjustment.

*Plasticity and Rehabilitation Indices*: We computed a “plasticity index” by dividing normalized projection density in uPyX-Rehab mice and uPyX+Rehab mice in a given subregion by the average Sham control value. This was correlated with a “rehabilitation index”, a proxy of rehabilitation-induced c-Fos activation in given subregions, which was calculated by dividing the average c-Fos density in runners divided by non-runners. Mice were the same as used for co-localization analyses. Only spinally-projecting regions (Wang et al., 2022) were included for these analyses.

*Statistical analyses*: Statistics were performed in R (version 4.4.1) using base R, lme4Test, and emmeans on blinded experimental groups. Footfalls, reach errors, and time to cross were calculated using custom-written code to analyze paw coordinate positions, as tracked by DeepLabCut. No main effects or interactions with sex were observed in any behavioral or anatomical measure, so data from males and females were pooled. For all statistical analyses, an alpha value of 0.05 was selected *a priori*, and p-values were adjusted for multiple comparisons.

## Results

### Voluntary rehabilitation engagement in uPyX and sham mice

We first asked whether mice with unilateral pyramidotomy (uPyX) would engage in complex-wheel running as a form of voluntary rehabilitation. After wheel habituation, mice received either a unilateral lesion of the left medullary pyramid or a sham surgery and were given access to runged wheels from 2 to 28 days post-injury (dpi) (**Fig. 1A**). Lesions were verified with PKC-γ immunostaining, and animals with incomplete lesions were excluded (**Fig. 1B-C**).

**Figure 1.**
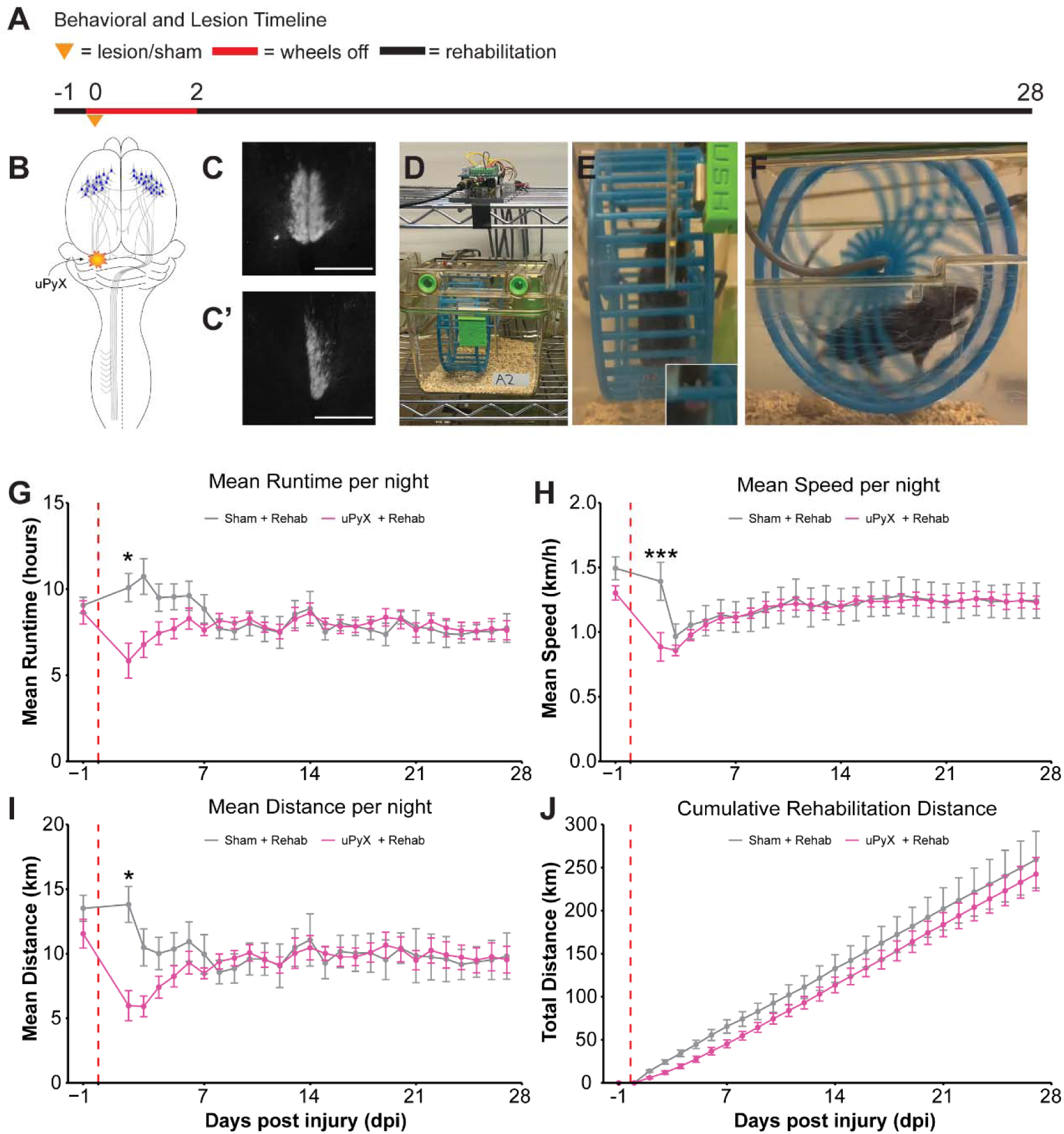
Mice with uPyX engage in significant voluntary complex wheel running rehabilitation. Mice had access to wheels for voluntary rehabilitation for 26 days post-lesion **(A).** Schematic of uPyX **(B)** and representative histological verifications of a sham and uPyX animals are shown **(C, C’)**. Wheel running data was collected using the REVS platform (**D**). Representative image of a mouse running, with inset showing forelimb rung grasping (**E**). Side-profile of mouse running **(F)**. Mean runtime from baseline to 28 dpi showed an acute deficit as revealed by an interaction effect between lesion and day (F_(2,374)_ = 5.295, p < 0.0001, ANOVA). Bonferroni-corrected post hoc testing revealed a significant difference at 2 dpi (p=0.01, t-test) **(G)**. Mean speed **(H)** and distance **(I)** also showed significant interaction effects (speed: (F_(2,374)_ = 4.469, p < 0.0001; distance: (F_(2,374)_ = 5.552, p < 0.0001; ANOVAs) and deficits at 2 dpi (speed: p = 0.03; distance: p = 0.00006; t-tests). Total cumulative distance ultimately showed no difference between lesioned and sham mice (F_(1,17)_=0.990, p=0.334, ANOVA) **(J)**. Dotted red line indicates lesion timepoint 0 dpi. Error bars represent +/− SEM. * = p < 0.01, *** = p < 0.001. Scale bar = 200 µm.

**Figure 1 Supplement 1.**
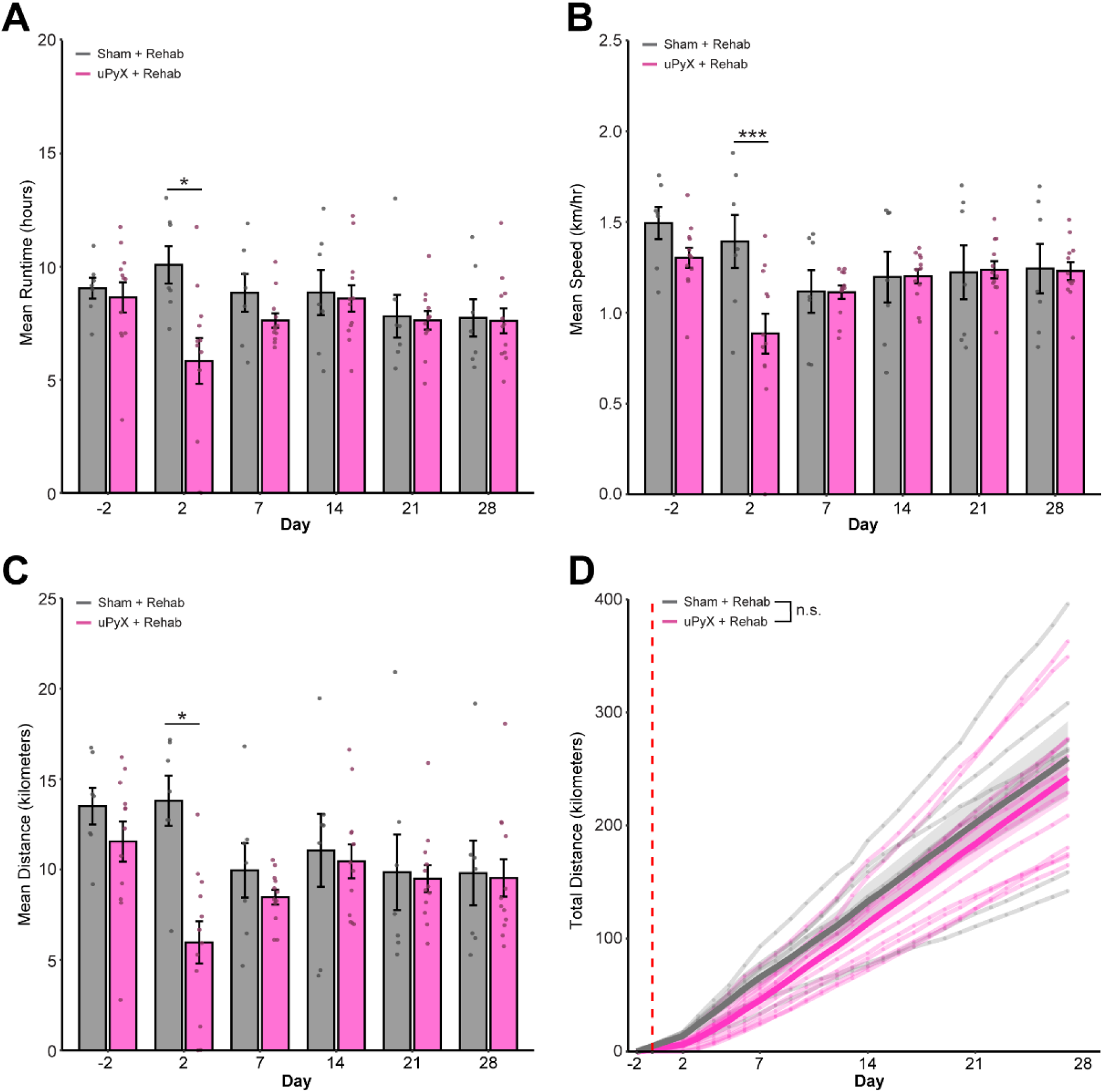
Rehabilitation metrics with individual data points. Bar plot showing mean runtime at – 2, 2, 7, 14, 21, and 28 dpi with individual data points. There was a significant deficit at 2 dpi (p=0.01, t-test) **(A)** between sham (gray) lesioned (pink) mice. Mean speed **(B)** and distance **(C)** also showed significant deficits at 2 dpi (speed: p = 0.03; distance: p = 0.00006; t-tests). Individual lines for each animal show total cumulative distance throughout the experiment for sham (gray) and lesioned (pink) mice. Thicker line shows group means and error bars represent +/− SEM. There was no significant effect of lesion (F(1,17)=0.990, p=0.334, ANOVA) **(D)**. Dotted red line indicates lesion timepoint, 0 dpi. * = p < 0.01, *** = p < 0.001.

Using the REVS platform (**Fig. 1D-F**) we acquired daily wheel running metrics. Across the 4-week rehabilitation period, uPyX and sham mice achieved comparable runtime, speed, and distance (**Fig. 1G-I**). There were no baseline differences observed between groups (p > 0.25 for all metrics), however, uPyX animals displayed a significant reduction in runtime, speed, and distance at 2 dpi (p < 0.01 for all comparisons), reflecting an acute motor deficit. These impairments resolved rapidly, with performance returning to sham levels by one week of rehabilitation (**Fig. 1G–I**, individual data points shown in **Fig. 1 – Supplement 1A-C**). Mixed-factor ANOVAs confirmed significant time × injury interactions for all running metrics (p < 0.001), but no long-term effect of injury on cumulative running (p = 0.33, **Fig. 1J**, individual animal line plots shown in **Fig. 1 – Supplement 1D**). Overall, these data show that uPyX mice fully re-engaged in voluntary complex-wheel running, enabling equivalent total rehabilitation between sham and lesioned mice.

### Rehabilitation enhances recovery of skilled forelimb function

To determine whether voluntary rehabilitation improved skilled motor performance, mice were tested on a complex ladder-rung task at baseline and at 2, 7, 14, 21, and 28 dpi (**Fig. 2A–B**). In addition to the uPyX+Rehab and Sham+Rehab groups, non-rehabilitated uPyX and Sham controls were included to isolate the effects of rehabilitation.

**Figure 2.**
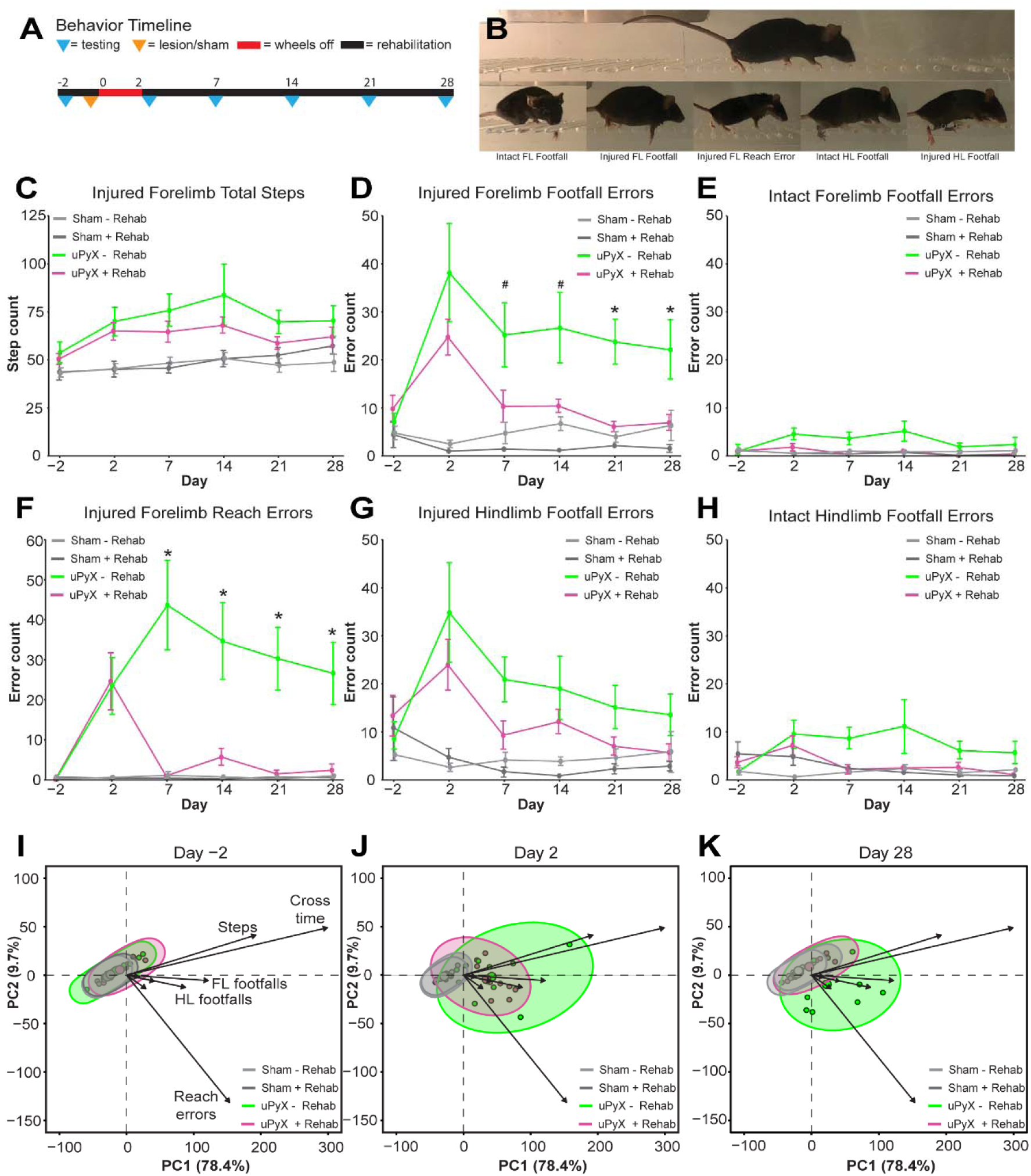
Rehabilitation promotes forelimb fine motor function recovery in uPyX mice. Complex ladder rung testing was completed on –2, 2, 7, 14, 21, and 28 dpi **(A)**. Representative image of a mouse crossing the complex ladder with examples of quantified errors **(B)**. Total number of injured forelimb (FL) steps significantly increased in lesioned animals (F_(1,33)_ = 14.445, p = 0.0006, ANOVA). This lesion effect persisted through 28 dpi (F_(1,32)_ = 13.339, p = 0.0009, ANOVA), but there was no effect of rehabilitation (F_(1,32)_ = 0.772, p = 0.402, ANOVA) (**C)**. Injured FL footfalls significantly increased in lesioned animals (F_(1,33)_ = 20.782, p = 0.00007, ANOVA) and this lesion effect persisted through 28 dpi (F_(1,32)_ = 12.090, p = 0.001, ANOVA). There was additionally a significant effect of rehabilitation (F_(1,32)_ = 7.938 p = 0.008, ANOVA). Bonferroni-corrected post hoc tests revealed a trend of uPyX+Rehab having improved performance relative to uPyX-Rehab at 7 dpi (p = 0.071, t-test) and 14 dpi (p = 0.052, t-test). There was a significant difference between the two groups at 21 dpi (p = 0.003, t-test) and 28 dpi (p = 0.035, t-test) (**D)**. The intact FL also showed a significant lesion-induced footfall deficit (F_(1,33)_ = 4.719, p = 0.037, ANOVA) at 2 dpi, but it did not persist through 28 dpi (F_(1,32)_ =2.186, p = 0.149, ANOVA). Although there was no long-term lesion effect, there was a main effect of rehabilitation (F_(1,32)_ = 4.37, p = 0.045, ANOVA) **(E)**. The injured FL had a significant increase in reach errors at 2 dpi (F_(1,33)_ =15.125, p = 0.00046, ANOVA). This effect persisted through 28 dpi (F_(1,32)_ =11.483, p = 0.002, ANOVA), and there was additionally a significant effect of rehabilitation (F_(1,32)_ = 9.171, p = 0.005, ANOVA). Bonferroni-corrected post hoc tests revealed that uPyX+Rehab significantly improved performance relative to uPyX-Rehab for injured FL reaches at 7 dpi (p = 0.00178, t-test), 14 dpi (p = 0.0114, t-test), 21 dpi (p = 0.00296, t-test), and 28 dpi (p = 0.00943, t-test) **(F)**. Injured HL footfalls had a significant lesion-induced increase at 2 dpi (F_(1,33)_ = 10.603, p = 0.003, ANOVA), which persisted through 28 dpi (F_(1,32)_ = 9.558, p = 0.004, ANOVA). There was a trend of rehabilitation improving injured HL footfalls (F_(1,32)_ = 3.347, p = 0.077, ANOVA) **(G)**. Intact HL footfalls significantly increased with lesion at 2 dpi (F_(1,33)_ = 4.317, p = 0.046, ANOVA) and this effect persisted through 28 dpi (F_(1,33)_ = 4.506, p = 0.042, ANOVA). There was a trend of rehabilitation improving intact HL footfalls (F_(1,32)_ = 3.510, p = 0.070, ANOVA) **(H)**. Principal component analyses revealed that all 4 groups clustered together at –2 dpi **(I)**. At 2 dpi, the two lesion groups separated from the sham groups, driven most strongly by differences in time to cross, step count, and injured FL errors **(J)**. By 28 dpi, the uPyX+Rehab separated from uPyX-Rehab and re-clustered with the sham groups **(K)**. Error bars represent +/−SEM. * = p < 0.05, = p<0.10.

At 2 dpi, uPyX animals exhibited significant increases in injured forelimb (FL) steps, injured FL footfalls and intact FL footfalls compared with sham controls (p < 0.05 for all, ANOVA), confirming a lesion-induced deficit (**Fig. 2C-E**). There was additionally a significant increase in injured FL reach errors (p < 0.001, ANOVA) (**Fig. 2F**). Across 7–28 dpi, rehabilitation significantly reduced footfall and reach errors. Both footfall and reaching errors showed main effects of lesion (footfall: p = 0.001; reach: p = 0.002, ANOVA) and rehabilitation (p = 0.008; reach: p = 0.005), with post hoc tests revealing that uPyX+Rehab mice improved progressively and reached near-sham levels by 21–28 dpi (p < 0.05 vs. uPyX-Rehab). The rehabilitation effect on reaching errors emerged as early as 7 dpi (p < 0.05 vs. uPyX-Rehab) and persisted throughout the testing period.

The injured and intact hindlimb (HL) also showed lesion effects (p < 0.05 for both, ANOVA) but were unaffected by rehabilitation (**Fig. 2G-H**). Individual data points across all days for the total steps, reach errors, and footfall errors are shown in **Fig. 2 – Supplement 2A-F**.

To visualize group-level behavioral patterns, we performed a principal component analysis (PCA) across all seven metrics (**Fig. 2I-K**). At baseline, all groups clustered together; at 2 dpi, both uPyX groups separated from shams. By 28 dpi, uPyX+Rehab mice re-clustered with sham controls, whereas uPyX-Rehab animals remained isolated (**Fig. 2K**).

To further investigate injured forelimb footfall and reach error severity on a finer, kinematic scale, we next plotted all injured forelimb footfalls and reaches with means at –2, 2, and 28 dpi **(Fig. 2 – Supplement 2A-F)**. There was little variation in displacement beneath the rung across groups and days for a typical forelimb footfall, suggesting that there is no effect of lesion or rehabilitation on severity of a missed step. By contrast, a typical reach was strongly overshot in all lesioned animals beginning on 2 dpi but resolved to baseline in rehabilitated animals only by 28 dpi. Together, these findings show that voluntary rehabilitation promotes robust and lasting recovery of fine forelimb control after unilateral CST injury.

**Figure 2 Supplement 1.**
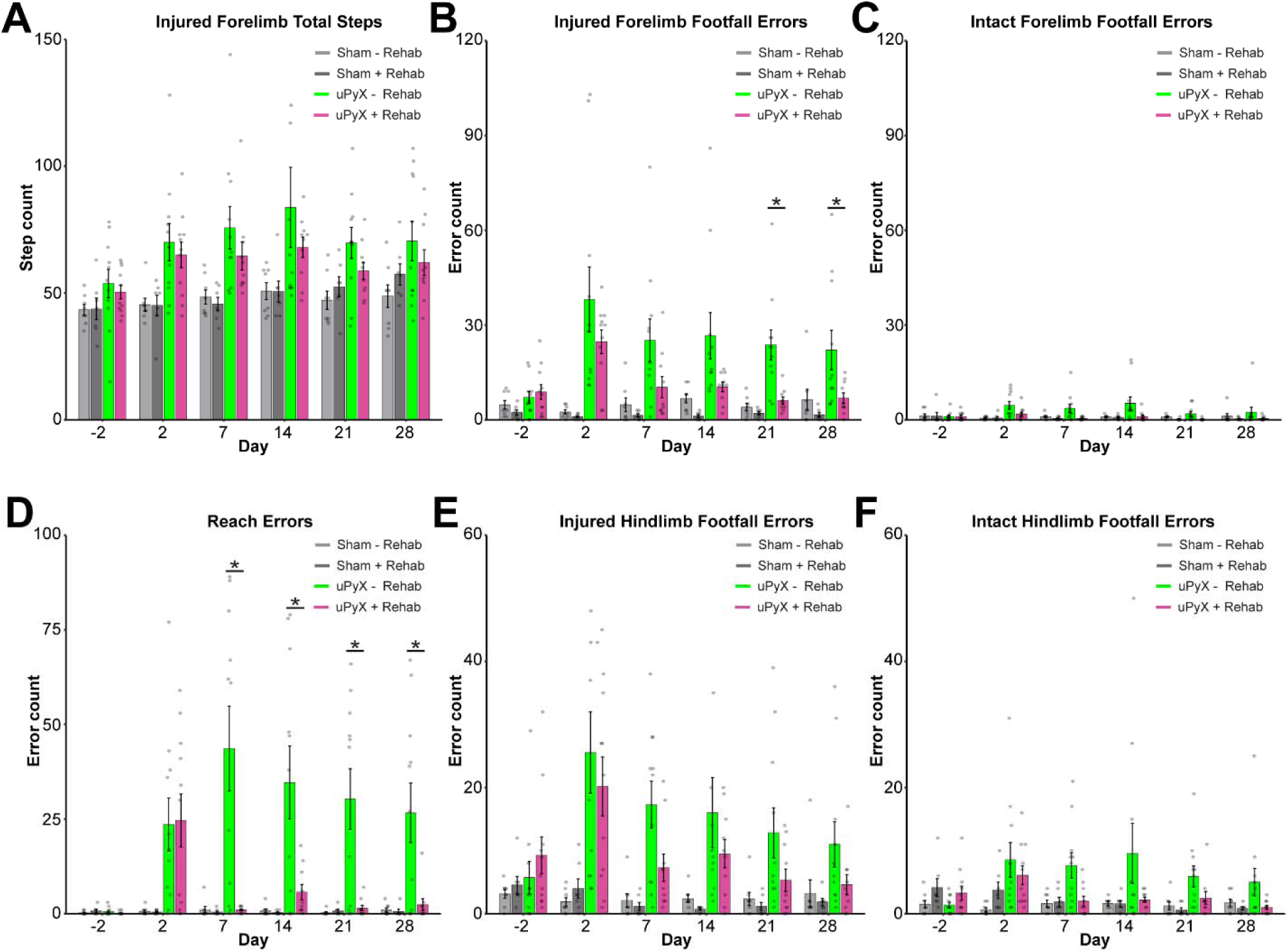
Fine motor function metrics with individual data points. Bar graphs at –2, 2, 7, 14, 21, and 28 dpi showing the total number of injured forelimb (FL) steps with individual data points. (**A)**. Bar graphs showed that injured FL footfalls significantly increased in lesioned animals (F_(1,33)_ = 20.782, p = 0.00007, ANOVA) and this lesion effect persisted through 28 dpi (F_(1,32)_ = 12.090, p = 0.001, ANOVA). There was additionally a significant effect of rehabilitation (F_(1,32)_ = 7.938 p = 0.008, ANOVA). Bonferroni-corrected post hoc tests revealed a trend of uPyX+Rehab having improved performance relative to uPyX-Rehab at 7 dpi (p = 0.071, t-test) and 14 dpi (p = 0.052, t-test). There was a significant difference between the two groups at 21 dpi (p = 0.003, t-test) and 28 dpi (p = 0.035, t-test) **(B)**. Bar graphs showing the total number of intact forelimb (FL) errors with individual data points **(C).** Bar graphs showing that the injured FL had a significant increase in reach errors at 2 dpi (F_(1,33)_ =15.125, p = 0.00046, ANOVA). This effect persisted through 28 dpi (F_(1,32)_ =11.483, p = 0.002, ANOVA), and there was additionally a significant effect of rehabilitation (F_(1,32)_ = 9.171, p = 0.005, ANOVA). Bonferroni-corrected post hoc tests revealed that uPyX+Rehab significantly improved performance relative to uPyX-Rehab for injured FL reaches at 7 dpi (p = 0.00178, t-test), 14 dpi (p = 0.0114, t-test), 21 dpi (p = 0.00296, t-test), and 28 dpi (p = 0.00943, t-test) **(D).** Bar graphs showing the total number of injured hindlimb (HL) errors with individual data points **(E)**. Bar graphs showing the total number of intact HL errors with individual data points **(F)**.

**Figure 2 Supplement 2.**
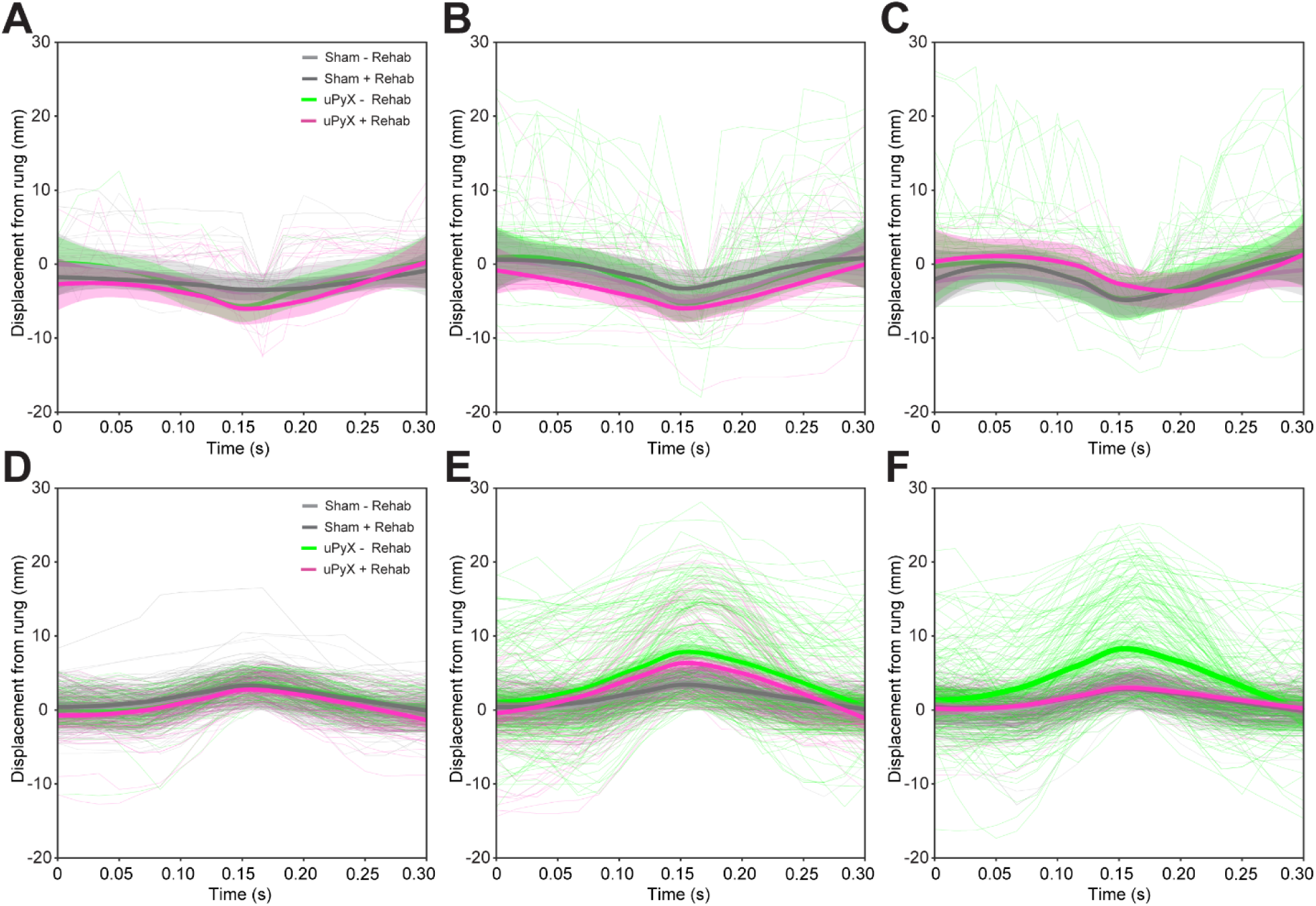
Individual and mean paw trajectories. Pooled individual injured forelimb footfalls from Sham-Rehab (light grey), Sham+Rehab (dark grey), uPyX-Rehab (green) and uPyX+Rehab (pink) groups at -2 dpi **(A)**, 2 dpi **(B)**, and 28 dpi **(C)** show consistent maximum displacement beneath the rung across days and groups. Thicker lines represent the mean typical footfall and shading represents the 99% confidence interval around the mean. Pooled individual injured forelimb reaches show typical reach height at -2 dpi **(D)**, aberrant overreaches in the lesioned groups at 2 dpi **(E)**, and recovery to baseline for rehabilitated mice at 28 dpi **(F)**. Thicker lines represent the mean typical reach and shading represents the 99% confidence interval around the mean.

### Complex-wheel running recruits intact corticospinal neurons

We next asked whether voluntary complex-wheel rehabilitation engaged spared CSNs. Using intersectional viral tracing, we labeled CSNs originating from the rostral (RFA) and caudal (CFA) forelimb motor areas in motor cortex and assessed neuronal activation with c-Fos immunostaining (**Fig 3A**). Mice were euthanized during their active cycle, and rehabilitation animals were confirmed to have run within the previous hour using REVS.

**Figure 3.**
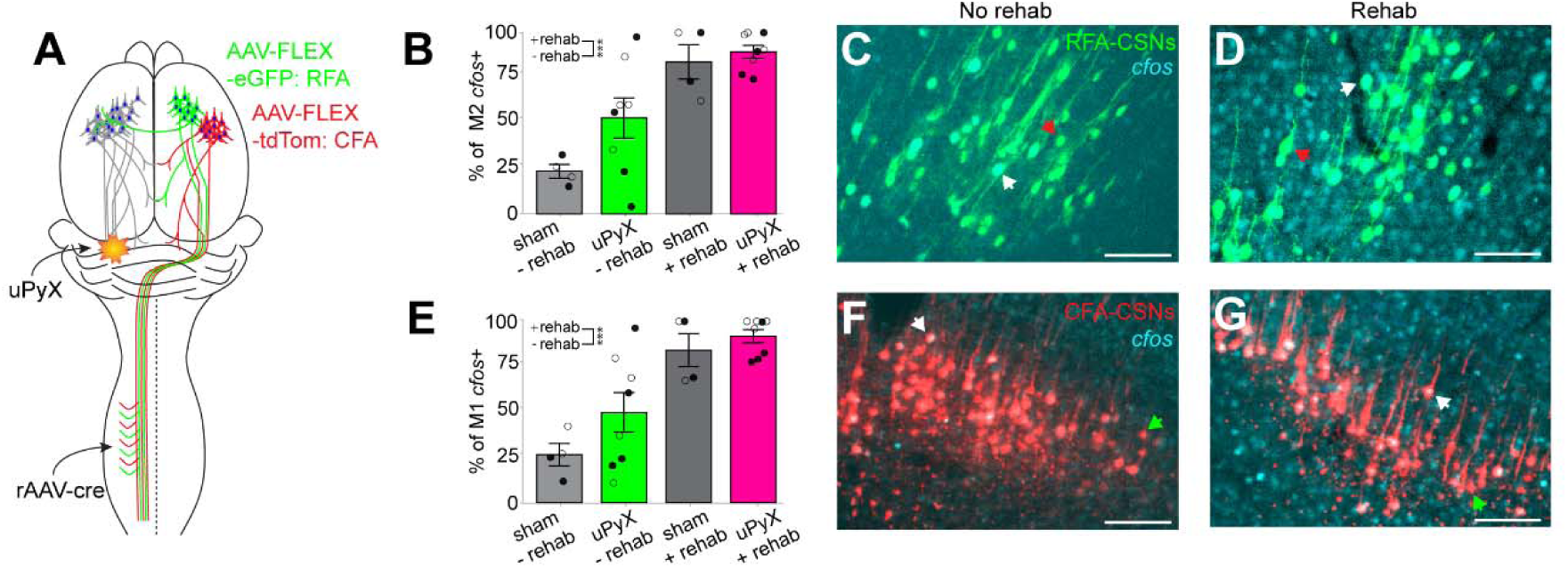
Rehabilitation functionally recruits intact CSNs. Schematic of intersectional tracing approach to target CSNs for labeling in M2 and M1 of mouse cortex **(A)**. There was a significant main effect of rehabilitation on percentage of cFos+ CSNs in M2, with rehab groups showing an increased colocalization (p=0.00538, linear mixed-effect model). Filled and open circles represent values from CFA– and RFA-traced cells respectively **(B)**. Representative images of M2 CSN and c-Fos labeling. Red arrows indicate a c-Fos-CSN and white arrows indicate a c-Fos+ CSN **(C, D)**. There was a significant main effect of rehabilitation on percentage of c-Fos+ CSNs in M1, with rehab groups showing increased colocalization (p=0.00389, linear mixed-effect model). Filled and open circles represent values from CFA– and RFA-traced cells respectively **(E)**. There was no effect of tracing origin on c-fos activation for both M2 (p=0.142, linear mixed-effect model) and M1 (p=0.079, linear mixed-effect model). Representative images of M1 CSN and c-Fos labeling. Green arrows indicate a c-Fos-CSN and white arrows indicate a c-Fos+ CSN **(F, G)**. Scale bar = 100 microns. Error bars represent +/− SEM. *** = p < 0.001.

CSNs localized to M1 and M2 using the Kim Atlas showed strong c-Fos expression following complex-wheel running, irrespective of their tracing origin, indicating robust recruitment of intact forelimb CSNs during rehabilitation (**Fig. 3B-G**). Across groups, there was a main effect of rehabilitation (M2: p = 0.005, M1: p = 0.004), but not lesion (M2: p = 0.21; M1: p = 0.26). This engagement suggested that the complex-wheel paradigm not only promotes behavioral recovery but also drives activity in spared CSNs.

### Rehabilitation drives CST-supraspinal terminal plasticity

Given the recruitment of intact CSNs, we next investigated whether rehabilitation induced supraspinal structural plasticity. Using the same intersectional labeling, we performed whole-brain projection density mapping with BrainJ to quantify CSN projections in 15 functional subregions spanning the cerebrum (blue), interbrain (green), midbrain (purple), pons (orange), medulla (pink), and cerebellum (yellow) (**Fig. 4A**).

**Figure 4.**
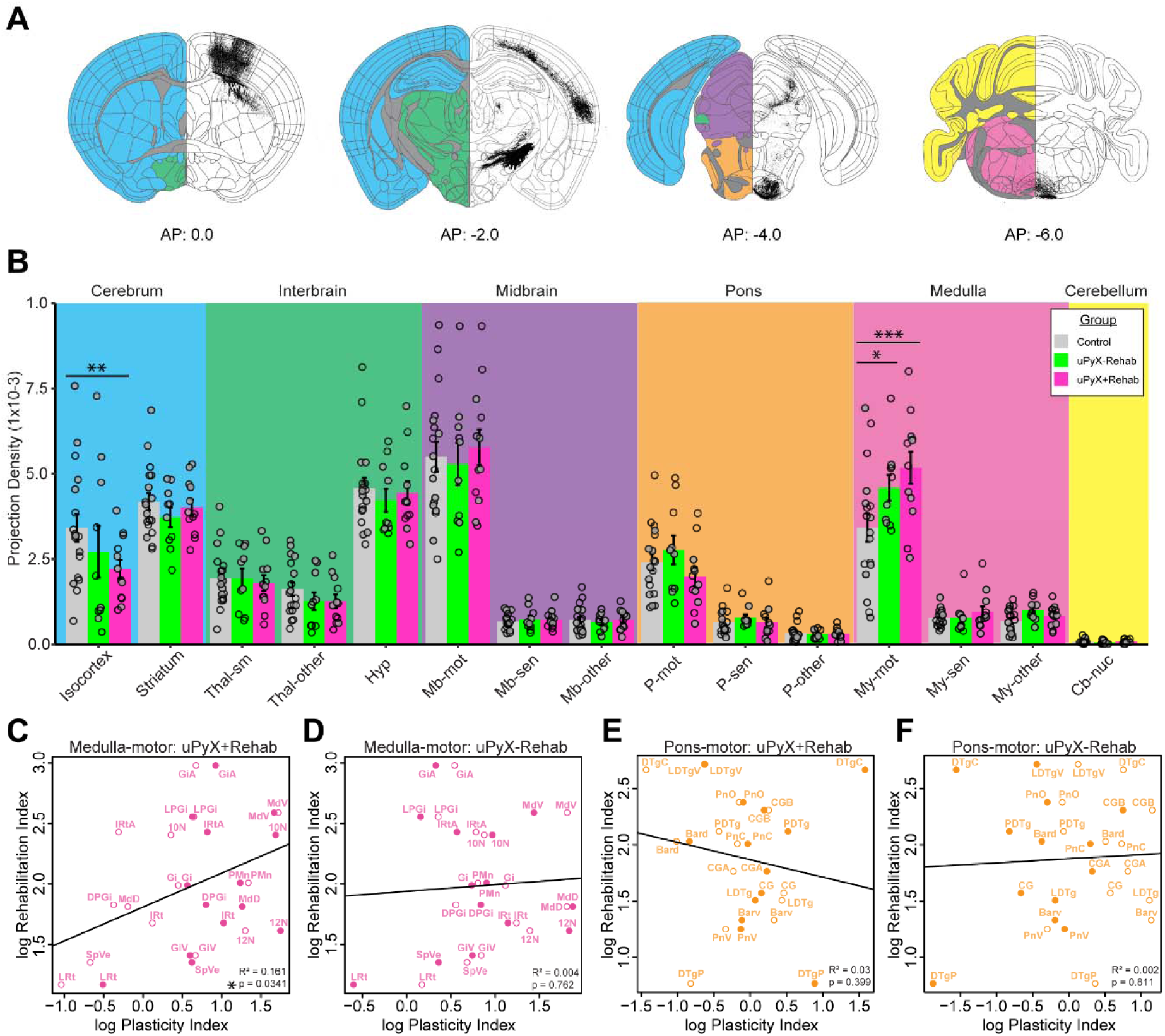
Large-scale reorganization of intact corticospinal tract projections in the lesioned and rehabilitated mouse brain. Representative intact corticospinal tract projections at 0.0 AP, –2.0 AP, –4.0 AP, and –6.0 AP from bregma. High-level organization of mouse brain regions are color-coded: blue = cerebrum, interbrain = green, midbrain = purple, pons = orange, medulla = pink, cerebellum = yellow. Traced projections are shown in black **(A)**. Quantification of normalized projection density in 15 functional subregions in control, uPyX-Rehab, and uPyX+Rehab mice. Closed and open circles represent values from CFA– and RFA-traced CSNs respectively. There was no significant effect of tracing origin on projection density (p=0.2447, linear mixed-effect model). The uPyX+Rehab mice had reduced projection density in the non-motor isocortex compared to control animals (p = 0.005587, linear mixed-effect model). uPyX-Rehab and uPyX+Rehab had increased projection density in the motor medulla compared to control animals (uPyX-Rehab vs. control, p = 0.0129; uPyX+Rehab vs. control, p = 0.000031; linear mixed-effect models). There were no significant differences in the other regions. **(B)**. Correlation between rehabilitation index and plasticity index for spinally-projecting medulla-motor regions in uPyX+Rehab and uPyX-Rehab mice. There was a significant correlation in uPyX+Rehab mice (R^2^ = 0.161, p = 0.0341, Pearson’s correlation), but not uPyX-Rehab mice (R^2^ = 0.004, p = 0.762, Pearson’s correlation) **(C, D)**. In spinally-projecting pons-motor regions, there were no significant correlations in uPyX+Rehab (R^2^ = 0.03, p = 0.399, Pearson’s correlation), nor uPyX-Rehab animals (R^2^ = 0.002, p = 0.811, Pearson’s correlation). Filled and open circles represent values from CFA– and RFA-traced cells respectively **(E, F)**. Error bars represent +/− SEM. * = p < 0.05, ** = p < 0.01, *** = p <0.001. Thal-sm=Sensory-motor thalamus, Thal-other=Thalamus other, Hyp=Hypothalamus, Mb-mot=Motor midbrain, Mb-sen=Sensory midbrain, Mb-other=Midbrain other, P-mot=Motor pons, P-sen=Sensory pons, P-other=Pons other, My-mot=Motor medulla, My-sen=Sensory medulla, My-other=medulla other, Cb-nuc=Cerebellar nuclei.

Projection density patterns were not significantly different between sham animals with or without rehabilitation; therefore, these groups were pooled as controls. Compared with controls, both lesion groups exhibited increased projection density in the motor regions of the medulla (uPyX+Rehab vs. Control, p < 0.001; uPyX-Rehab vs. Control, p = 0.013), with the largest changes observed in rehabilitated animals (**Fig 4B**). In contrast, non-sensory/motor regions of the isocortex showed decreased CST projection density in uPyX+Rehab mice (uPyX+Rehab vs. Control, p = 0.005). Thus, rehabilitation induced targeted rather than global CST-supraspinal reorganization, with pronounced effects in brainstem motor nuclei. All other subregions showed no significant difference between groups. Additionally, there was no significant effect of tracing origin on CST projection density in downstream regions.

To link neural engagement during rehabilitation to plasticity of intact CST projections, we derived a “rehabilitation index” from regional c-Fos activation and a “plasticity index” from normalized CST projection density relative to controls. Across spinally projecting regions, these indices were significantly correlated in the motor medulla of uPyX+Rehab mice but not in uPyX-Rehab animals (Pearson’s correlation; R^2^ = 0.161, p = 0.031), indicating that CST reorganization occurs preferentially in regions most activated during rehabilitation (**Fig. 4C–D**). No such correlation was observed in a control region (motor pons), suggesting region-specific, activity-dependent plasticity (**Fig. 4E–F**).

### Rehabilitation-induced plasticity localizes to specific motor medulla nuclei

Because the motor medulla emerged as a major locus of CST reorganization, we next examined projection density at the level of individual medullary nuclei. Mixed-effects models identified three nuclei showing significant lesion– and/or rehabilitation-induced plasticity: the lateral paragigantocellular reticular nucleus (LPGi), the alpha part of the gigantoreticular nucleus (GiA), and the ventral medullary reticular nucleus (MdV) (**Fig. 5A-C).** Projection density increased in the LPGi (p = 0.010) and GiA (p < 0.001) specifically in uPyX+Rehab animals relative to controls, whereas the MdV exhibited increases in both uPyX-Rehab (p = 0.004) and uPyX+Rehab (p < 0.001) groups, suggesting contributions from both lesion and rehabilitation (**Fig. 5A–D**). For all three of these subregions, there was no significant effect of tracing origin on CST projection density. Representative images show intact CST projection density in the rostral (**Figs. 5D-G**) and caudal (**Figs. 5H-K**) medulla.

**Figure 5.**
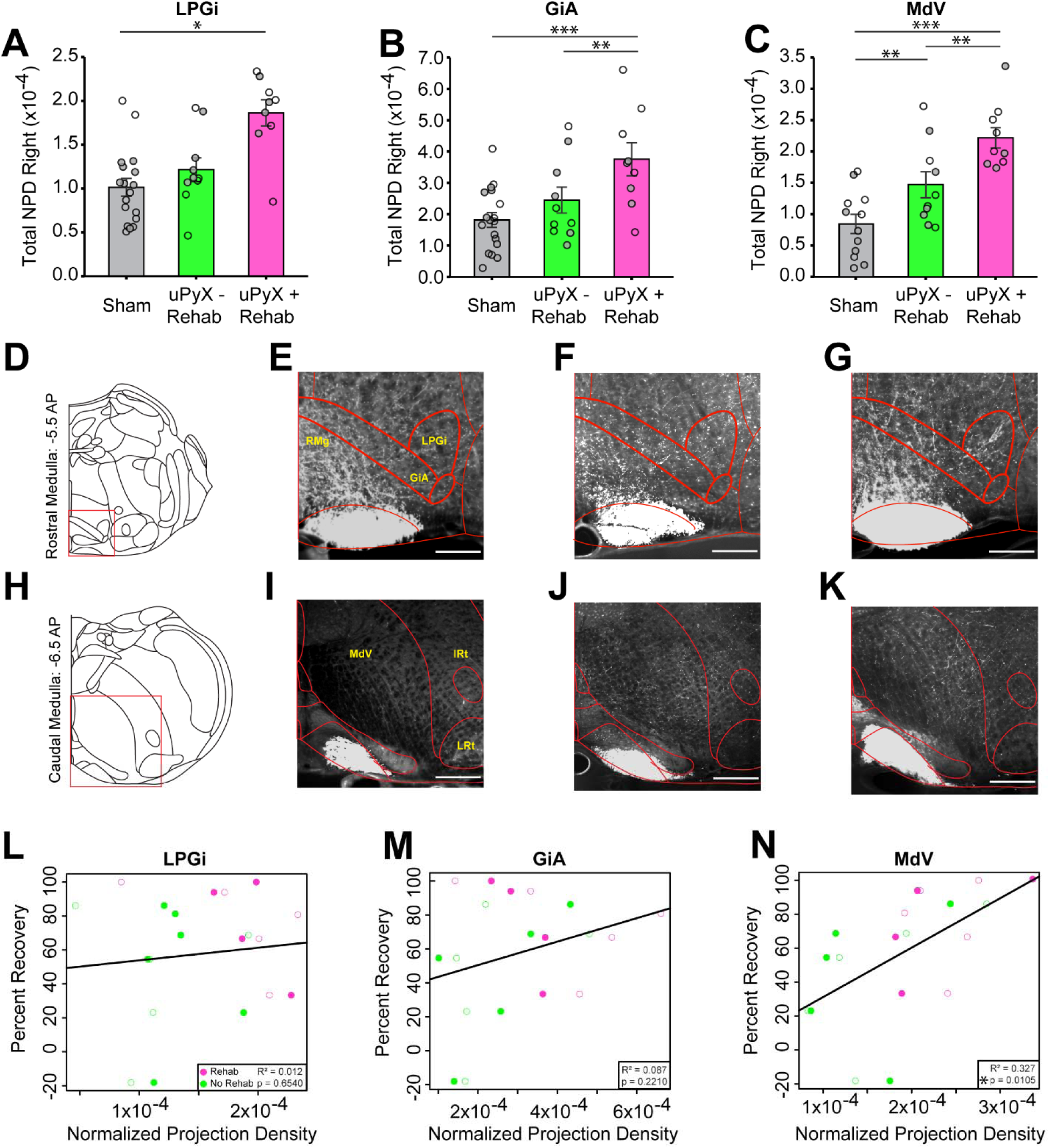
Subregion specific reorganization of intact corticospinal tract projections in the medulla of lesioned and rehabilitated mice. Normalized projection density (NPD) increased in the lateral paragigantoreticular nucleus (LPGi) of uPyX+Rehab mice compared to control mice (t=2.91, p=0.01, Tukey-adjusted post hoc). There was no main effect of tracing origin (t=0.850, p=0.408; linear mixed-effect model) **(A)**. Normalized projection density increased in both uPyX+Rehab (t = 6.67, p < 0.0001, Tukey-adjusted post hoc) mice compared to controls, and uPyX+Rehab mice compared to uPyX-Rehab mice (t = 3.98, p = 0.002, Tukey-adjusted t-test) in the alpha part of the gigantoreticular nucleus (GiA). There was no main effect of tracing origin (t=0.851, p=0.407; linear mixed-effect model) **(B)**. Normalized projection density increased in both uPyX+Rehab (t = 7.11, p < 0.0001, Tukey-adjusted post hoc) mice compared to controls, and uPyX+Rehab mice compared to uPyX-Rehab mice (t = 3.70, p = 0.001706, Tukey-adjusted t-test) in the ventral part of the medullary reticular nucleus (MdV). Normalized projection density also increased in uPyX-Rehab mice compared to controls (t = 3.34, p = 0.004967, Tukey-adjusted t-test). There was no main effect of tracing origin (t=1.493, p=0.157; linear mixed-effect model) **(C)**. Schematic of the rostral medulla (**D**) and representative images of intact CST density in a control **(E)**, uPyX-Rehab **(F)**, and uPyX+Rehab **(G)** mouse. Schematic of the caudal medulla **(H)** and representative images of intact CST density in a control **(I)**, uPyX-Rehab **(J)**, and uPyX+Rehab **(K)** mouse. The red box indicates the zoomed-in area of interest. Correlations between percentage of injured forelimb footfall recovery and normalized projection density were not significant in the LPGi (R^2^ = 0.012, p = 0.654, Pearson’s correlation) nor GiA (R^2^ = 0.087, p = 0.221, Pearson’s correlation) **(L, M)**. Correlation between percentage of injured forelimb footfall recovery and normalized projection density was significant in the MdV (R^2^ = 0.327, p = 0.0105, Pearson’s correlation). Closed and open circles represent values from CFA– and RFA-traced CSNs respectively **(N)**. Error bars represent +/− SEM. * = p < 0.05, ** = p < 0.01, *** = p < 0.001. Scale bar = 200 microns.

Crucially, to determine the functional relevance of these changes, we correlated CST projection density in each nucleus with behavioral recovery. Projection density in the MdV, but not LPGi or GiA, significantly correlated with improved forelimb footfall recovery (Pearson’s correlation; R^2^ = 0.327, p = 0.011; **Figs. 5D–F**). These findings reveal the MdV as a key nucleus where intact CST structural plasticity is directly linked to functional improvement.

## Discussion

Here, we show that voluntary, fine motor-dependent rehabilitation drives robust recovery of skilled forelimb function after partial CST injury and induces highly targeted CST-supraspinal terminal plasticity. By combining uPyX with continuous complex-wheel training and whole-brain projection mapping, we identify specific medullary reticular nuclei that undergo rehabilitation-dependent CST remodeling and link this structural plasticity to functional activation and behavioral recovery. These results provide new insight into how voluntary motor rehabilitation engages and reorganizes intact supraspinal pathways to support functional recovery after SCI.

### Voluntary rehabilitation drives recovery of skilled motor function

Recovery of motor function after SCI is dependent in part on the capacity of spared, descending tracts to reorganize and reintegrate with impaired motor circuits (Fink and Cafferty, 2016). Motor rehabilitation of the affected limbs is key to this process in humans (Lee and Jeoung, 2023) and rodent models (Mensah et al., 2025). However, the anatomical mechanisms through which rehabilitation promotes functional recovery are unknown. Despite significant damage to descending motor pathways, human SCIs are rarely anatomically complete (Burns et al., 2012), allowing residual motor circuits to serve as substrates for activity-dependent re-wiring via rehabilitation. This underscores the need for models that promote sustained motor engagement after SCI.

Here, we established a voluntary rehabilitation model that maintains high levels of motor activity after partial CST injury. The unilateral pyramidotomy model (uPyX) results in a specific unilateral lesion to the CST while preserving other sensorimotor pathways. This allows animals to retain gross locomotor function yet display reproducible fine motor deficits (Starkey et al., 2011). Thus, the uPyX model is well suited for studying activity-dependent recovery of fine motor function and plasticity in spared corticospinal circuits.

Using our REVS platform (Bonanno et al., 2025) for continuous wheel-running monitoring, we confirmed that uPyX mice engaged in sustained, self-directed wheel running throughout the rehabilitation period (**Fig. 1**). To selectively engage the CST for fine motor control (Iwaniuk and Whishaw, 2000), we used complex rung-patterned wheels. Consistent with previous work showing that lesioned animals can attain sham levels of wheel running shortly after injury (Loy et al., 2018), we found that uPyX mice quickly restored their capacity for complex, voluntary wheel running (**Fig. 1**). This indicates that spared CST circuits remain highly recruitable after uPyX, providing a foundation for recovery of skilled motor function and rehabilitation-induced plasticity. Indeed, voluntary rehabilitation led to significant improvement of functional forelimb metrics, with rehabilitated mice achieving sham-level accuracy on the complex ladder rung task within 1-3 weeks post-injury (**Fig. 2**). Recovery of forelimb footfalls and forelimb reach errors is consistent with the restoration of typical corticospinal control and suggests that rehabilitation led to more than just gross locomotor improvements.

Notably, prior studies show that reach training rehabilitation paradigms induce task-specific recovery but impair performance on untrained tasks including the complex ladder rung task (Girgis et al., 2007; Krajacic et al., 2010). By contrast, voluntary complex-wheel running here led to task-non-specific improvements that generalized to the complex ladder rung task. This suggests that high levels of CST-targeted voluntary motor training may facilitate more general restoration of forelimb function. Overall, these findings validate our model as suitable for studying rehabilitation-induced functional recovery after partial SCI and motivate an investigation of how voluntary rehabilitation recruits spared corticospinal pathways.

### Mechanisms of Voluntary Rehabilitation-Driven Plasticity in Spared Corticospinal Circuits

We found that voluntary complex-wheel running increased c-Fos expression in intact CSNs from M1 and M2 of forelimb motor cortex (**Fig. 3**), confirming that rehabilitation engages intact CST pathways which are critical for skilled motor control after injury (Carmel et al., 2010; Wahl et al., 2014). Moreover, this neural recruitment can facilitate activity-dependent plasticity in spared CSN circuits, which have previously been described to functionally reorganize after injury and rehabilitation (Weidner et al., 2001; Dancause et al., 2005; Ghosh et al., 2009).

Whole-brain projection mapping revealed that rehabilitation-induced intact CST structural remodeling is highly targeted to specific supraspinal brain regions. Significant increases in CST projection density were localized to the motor-medulla, specifically the lateral paragigantocellular nucleus (LPGi), the alpha part of the gigantoreticular nucleus (GiA), and the ventral medullary reticular nucleus (MdV). We did not detect a difference in RFA– and CFA-traced projection density for these three regions, consistent with a previous report showing similar innervation patterns by RFA– and CFA-CSNs throughout the motor medulla (Carmona et al., 2024). In contrast, the non-sensorimotor isocortex showed reduced CST projection density after rehabilitation, suggesting a pruning or reorganization of projections away from less functionally relevant targets (Kleim and Jones, 2008).

Novelly, we found that the extent of CST plasticity within medullary subregions strongly correlated with the level of neuronal activation induced by rehabilitation in those same regions (**Fig. 4**). This supports a model of rehabilitation-induced region-specific functional reorganization, in which rehabilitation selectively strengthens intact projections to circuits most engaged by fine motor control. Consistent with this model, studies have demonstrated that CST collateral sprouting and synaptogenesis is dependent on post-synaptic partner activity and signaling (Ueno et al., 2012; Bradley et al., 2019).

The medullary reticular nuclei which underwent CST remodeling here have distinct, complementary roles in fine motor control (Inoue and Ueno, 2025). The MdV is critical for skilled forelimb function such as grasping (Esposito et al., 2014). In line with this functional role, intact CST projection density in this subregion correlated with forelimb footfall percentage recovery (**Fig. 5**). The LPGi and GiA primarily contribute to speed modulation and locomotion initiation (Capelli et al., 2017; Zhang et al., 2024). While we observed *de novo*, rehabilitation-specific increases in CST density in these regions, CST density here did not correlate with forelimb footfall recovery. Nevertheless, rehabilitation-induced sprouting in these regions may play a supporting role in reorganizing motor control networks to re-establish the skilled motor function observed in rehabilitated mice. Additionally, plasticity in these nuclei may play a more consequential role in functional recovery after more severe SCI (Lemieux et al., 2024).

The MdV, LPGi, and GiA project predominantly ipsilaterally to the spinal cord (Esposito et al., 2014; Liang et al., 2016), positioning these nuclei as potential conduits for conveying descending motor commands to the denervated side after unilateral injury. This ipsilateral organization provides a plausible anatomical substrate through which spared CST axons can reroute motor output via reticulospinal pathways to the denervated spinal circuits. Crucially, these reticulospinal circuits are often spared after contusion injury (Wang et al., 2022).

Previous work has highlighted the importance of plasticity within descending reticulospinal and midbrain pathways (Filli et al., 2014; Lemieux et al., 2024; Jeleva et al., 2026). Our findings extend this framework by demonstrating that rehabilitation also induces plasticity of corticospinal inputs onto these same brainstem nuclei. This finding aligns with the established role of the CST in modulating both supraspinal and spinal motor circuits in intact and injured states (Liu et al., 2018; Ueno et al., 2018; Moreno-Lopez et al., 2021; Glover and Baker, 2022). Moreover, after injury, CST axons have been shown to form detour circuits through propriospinal and brainstem relays (Bareyre et al., 2004; Asboth et al., 2018), providing support for the cortico-reticular remodeling observed here.

Our study builds upon the cortico-reticular model proposed by Asboth et al., 2018. Firstly, we show that CST projections, rather than intratelencephalically-projecting motor neuron projections, undergo terminal plasticity. Secondly, we identify the specific medullary subregions of the reticular formation that are modulated in rehabilitated mice. Finally, we demonstrate that there are unique patterns of sprouting amongst these different reticulospinal sub-regions (i.e, MdV experienced uPyX-Rehab and uPyX+Rehab sprouting, while LPGi/GiA only experienced uPyX+Rehab sprouting).

Together, our findings support a model in which rehabilitation selectively promotes CST sprouting toward medullary nuclei that are anatomically and functionally poised to re-establish descending control of the impaired limb.

### Limitations and Future Directions

While our rehabilitation intervention was causally linked to improvements in motor behavior, anatomical analyses of the intact CST presented here are correlational and do not by themselves establish that CST sprouting within the motor medulla directly drives recovery. Nevertheless, we believe that the strong association between CST projection density in the MdV and functional improvement suggests a potential causal contribution that warrants further testing. Indeed, our previous data showed that chemogenetic silencing of lesion-induced sprouting of rubro-raphe circuits abrogated recovery of fine forelimb function after bilateral pyramidotomy (Siegel et al., 2015). The present work provides a foundation for similar studies using region-specific circuit manipulation to directly determine the role of corticospinal–medullary pathways in mediating rehabilitation-induced recovery across various SCI models.

A second consideration is the translational relevance of the uPyX model. Although it most closely resembles partial lesions such as Brown–Séquard syndrome and unilateral cortical stroke, its primary advantage is its ability to isolate the capacity for CST reorganization, which ultimately uncovered three specific medullary nuclei associated with rehabilitation-induced recovery. Finally, the molecular and cellular mechanisms that govern CST-supraspinal remodeling remain incompletely defined. Elucidating these mechanisms and determining how targeted rehabilitation paradigms might selectively engage homologous cortico-reticular systems in humans, represents a key step toward optimizing activity-based therapies to enhance functional recovery after SCI.

## Acknowledgements

This work was supported by the National Institutes of Health National Institute of Neurological Disorders and Stroke Grants R01NS121026 and R21NS139481, and Wings for Life 2023-066.

